# Ancestral dietary change alters development of *Drosophila* larvae through MAPK signalling

**DOI:** 10.1101/2020.10.07.327478

**Authors:** Samuel G. Towarnicki, Neil A. Youngson, Susan M. Corley, Jus C. St John, Nigel Turner, Margaret J. Morris, J. William O. Ballard

**Author notes:** Electronic addresses. Contact J. William O. Ballard School of Biotechnology and Biomolecular Sciences, The University of New South Wales, Sydney, Australia 2052.

## Abstract

Increasing evidence in animal species ranging from mammals to insects has revealed phenotypes that are caused by ancestral life experiences including stress and diet. The descendent phenotypes themselves are wide ranging, and include changes to behaviour, disease risk, metabolism, and growth. Ancestral dietary macronutrient composition, and quantity (over- and under-nutrition) have been shown to alter descendent growth, metabolism and behaviour. Several studies have identified inherited molecules in gametes which are altered by ancestral diet and are required for the transgenerational effect. However, there is less understanding of the developmental pathways in the period between fertilisation and adulthood that are altered by the inherited molecules. Here we identify a key role of the MAPK signalling pathway in mediating changes to *Drosophila* larval developmental timing due to variation in ancestral diet. We exposed grand-parental and great grand-parental generations to defined protein to carbohydrate (P:C) dietary ratios and measured developmental timing. Descendent developmental timing was consistently faster in the period between the embryonic and pupal stages when the ancestor had a higher P:C ratio diet. Transcriptional analysis of embryos, larvae and adults revealed extensive and long-lasting changes to the MAPK signalling pathway which controlled growth rate through regulation of ribosomal RNA transcription. The importance of these processes was supported by pharmacological inhibition of MAPK and rRNA proteins which reproduced the ancestral diet-induced developmental changes. This work provides insight into the role of developmental growth signalling networks in mediating non-genetic inheritance in the period between fertilisation and adult.

**Summary statement:** Ancestral, diet-induced descendent developmental timing changes are caused by alteration of MAPK signalling pathways in the period between the embryo and pupal stages in *Drosophila*.

## Introduction

Evidence accumulated across animal and plant research over the last 20 years has confirmed that the inherited determinants of an organism’s phenotype are more than just DNA (Cavalli and Heard, 2019). Inheritance of RNA, protein and metabolites in the male or female gamete can influence a variety of traits such as size, shape, behaviour and health (Legoff et al., 2019; Perez and Lehner, 2019). Furthermore, the levels and types of inherited molecules can be influenced by ancestral environmental exposures. The consequences of these molecules on descendent phenotype range from a proposed short-term mode of adaptation which contrasts with longer term DNA mutation and selection (Auge et al., 2017; Prokopuk et al., 2015), to contributing to an individual’s disease risk (Nilsson et al., 2018).

Ancestral environmental changes that have been found to alter phenotypes of descendants include behavioural stress, toxin exposure and nutritional variation (Cavalli and Heard, 2019; Legoff et al., 2019; Perez and Lehner, 2019). The latter is perhaps the most-studied, and examples of ancestral diet altering descendent phenotype have been documented due to over- or under-nutrition in natural populations including human (Aiken and Ozanne, 2014) and laboratory organisms such as *Caenorhabditis elegans* (Kishimoto et al., 2017; Tauffenberger and Parker, 2014), *Drosophila* (Emborski and Mikheyev, 2019; Ost et al., 2014; Strilbytska et al., 2020; Vijendravarma et al., 2010; Xia et al., 2016) and rodents (Aiken and Ozanne, 2014). In these studies changes to descendent metabolism and growth is often reported, resulting in developmental timing, organ size and body weight alterations. For example, in *Drosophila*, paternal sugar exposure reprogrammed offspring lipid metabolism leading to an increase in stored triglyceride levels (Ost et al., 2014). Over 100 studies in rodents have also shown reprogramming of offspring feeding behaviour, birth and adult body weight, adiposity, insulin/glucose metabolism, hypertension etc due to maternal or paternal obesity or starvation (Aiken and Ozanne, 2014). Another important feature of developmental programming is that the effect on descendants is dependent on the timepoint of exposure to the environmental change (Lecomte et al., 2013). Frequently effects are noted in offspring generation, but are absent, weaker or even different in the grand-offspring generation (Aiken and Ozanne, 2014). This is due to offspring always experiencing the exposure directly, either across the placenta while *in utero*, or as exposure to germ cells that will go on to become offspring. The transmission mechanisms can also be indirect, for example through faecal microbiota transfer, due to alterations in maternal care (including lactation). However, a major focus of the developmental programming field has been in the identification of molecules in the gametes that drive descendent phenotypes (Perez and Lehner, 2019). Epigenetic changes such as DNA methylation, histone modifications and non-coding RNAs have all been found to alter offspring gene expression resulting in developmental and adult phenotype differences. In the aforementioned *Drosophila* model, sugar exposure extensively altered repressive sperm histone methylation patterns in the exposed males, with fat metabolism genes being particularly affected and prone to subsequent transcriptional changes in offspring, but not grand-offspring (Ost et al., 2014). In mammals the physiological intimacy of gestation means that developmentally programmed offspring phenotypes are hard to prove to be due to gamete-derived molecules if the mother is the exposed parent. However, this is not the case when the father is the exposed parent and there are strong examples of paternal diet- and stress-induced offspring phenotypes being due to alterations in sperm small non-coding RNAs (Chen et al., 2016; Gapp et al., 2014; Klastrup et al., 2019).

In contrast to the abundance of studies that have searched for a causative epigenetic change in the gamete, there is less information on the post-fertilisation mechanisms that bridge the reading of the inherited molecule and the ultimate cell, tissue or whole organism phenotypes. As mentioned most developmental programming studies have reported metabolic phenotypes and work by ourselves and others has suggested that developmentally programmed changes to pancreas function in mammals are mediated by Foxo1-(Talchai and Accili, 2015) and Myc-directed pathways (Ng et al., 2014). However, less has been reported on changes to pathways that regulate offspring growth and developmental timing. Therefore, in this study we aimed to gain a deeper understanding of the developmental pathways that are affected by ancestral diet at different offspring life stages.

In our previous study we observed generational differences in the different *Drosophila* strains that were fed diets of varying protein to carbohydrate (P:C) ratios (Aw et al., 2018). We hypothesised that the generational differences could be due to transgenerational epigenetic inheritance effects that altered the rate at which the flies progress through the developmental stages of embryo, larva, pupae, and adult. Importantly, these effects appeared to be somewhat accumulated across generations suggesting that they were not solely caused by the diet that the fly was consuming when it produced its gametes (Strilbytska et al., 2020; Vijendravarma et al., 2010). We therefore designed our experiments to exclude parental effects by ensuring that all parents consumed the same diet, and variation in ancestral diet was limited to the grand-parental and great grand-parental generations (Figure 1). Our studies confirmed ancestral P:C diet-induced descendent developmental timing changes, which were due to alteration of MAPK signalling pathways in the period between the embryo and pupal stages.

**Figure 1.**
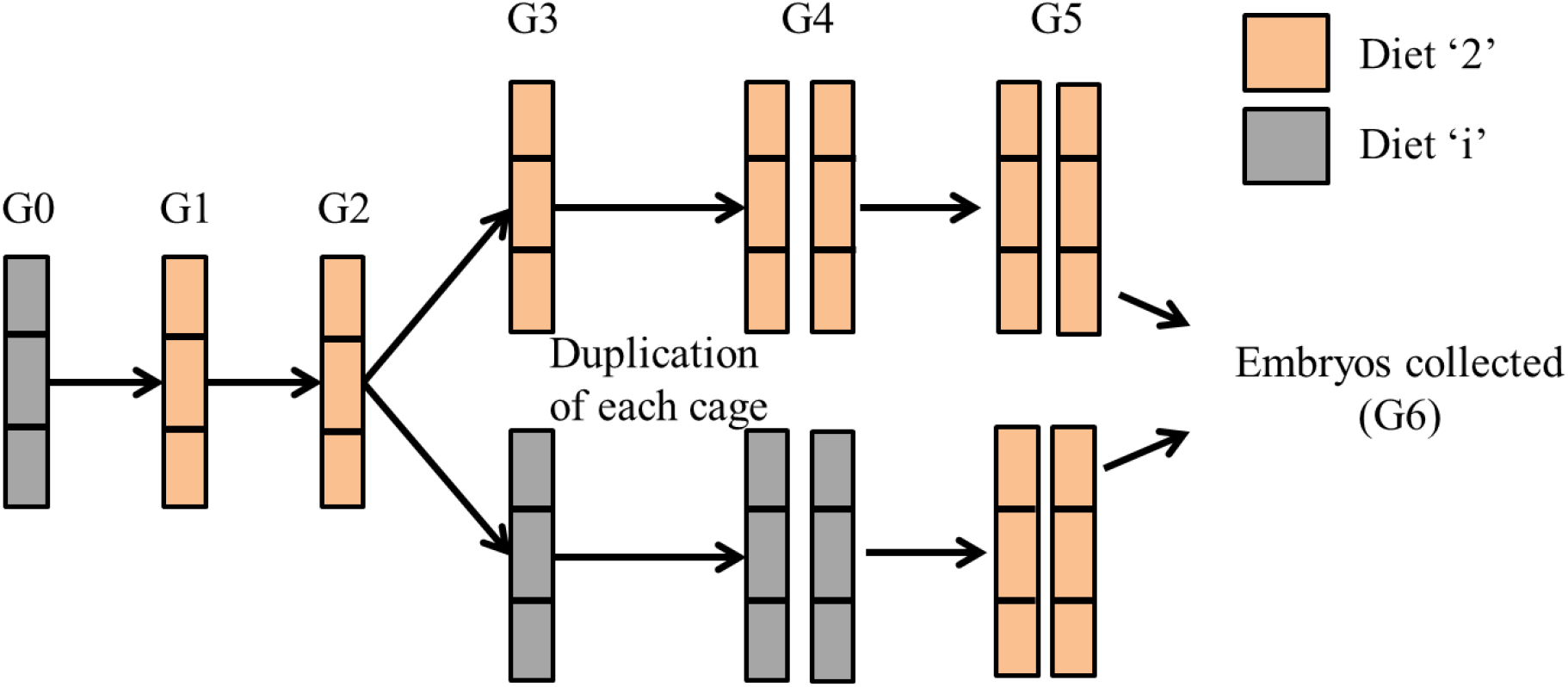
Overview of experimental design: The generational design allowed comparison of G6 embryos produced from great-grandparents and grandparents that differed in their dietary composition, with parents provided the same diet.

## Materials and methods

### Diets and fly genotypes

We utilise four diets in this study that differ in their P:C ratios. Three of these diets, 1:2, 1:4 and 1:6 P:C (referred to as diet ‘2’, diet ‘4’, and diet ‘6’, respectively) are constructed diets, and one, Formula 4-24 instant *Drosophila* medium (Carolina biological supply company, referred to as diet ‘i’) is a pre-made diet. We use the Alstonville;*w*^1118^ strain of flies used by Aw et al. (2018).

### Generational experimental design

We utilised a generational design (Fig. 1).

Initially a single population cage was established with ∼800 adult Alstonville flies (termed generation 0). Oviposition medium consisting of 10% treacle and 4% agar with a thin spread of baker’s yeast was placed into the cage. This medium was replaced daily for 3 d. On the third day flies were provided with the oviposition medium for 6 hours before the medium was removed, and embryos were washed and collected as per Clancy and Kennington (2001). Embryos were then placed into 12 bottles containing diet ‘2’ (generation 1) at ∼200 embryos in each bottle, with four bottles per population cage. For all further generations, when flies were observed to be eclosing, flies from that day were discarded, and counted as day 0. After a further 72 hours the bottles were opened, adult flies populated the cages, and all bottles were removed. Embryos were collected as described previously and established on the diet ‘2’ again (generation 2).

Embryos collected from generation 2 were then used to seed bottles with diets ‘i’, ‘2’, ‘4’ or ‘6’ diet (generation 3). Embryos from generation 3 were moved onto the same diet as their parents (i.e. ‘i’ to ‘i’, ‘2’ to ‘2’ etc.; generation 4), and the number of cages was doubled at this generation from 3 to 6 cages. Finally, embryos from generation 4 were placed onto bottles containing diet ‘2’ (generation 5).

Embryos produced by generation 5 were collected within 45 minutes of laying, and were used for RNA-seq, RT-qPCR, and development assays. Collected embryos (generation 6) are referred to as their previous 3 ancestral generations, those maintained consistently on diet ‘2’ are termed 222, while those fed different ancestral diets are termed ii2, 442, and 662.

Partial inhibition of the MAPK pathway and Ribosome biogenesis pathway was conducted through addition of inhibitors to diet ‘2’ for generation 6 embryos. For MAPK pathway *Egfr* was inhibited with 100 μM Tyrphostin AG1478, and *rolled* was inhibited using 2.5 μM SCH772984. Ribosome biogenesis was inhibited at *pol1* using 50 μM BMH-21.

### Development assays

Development of generation 6 embryos was conducted using diet ‘2’ and was measured as count of adult flies eclosing (hatching) in 3 d, and time to pupation. Time to pupation is measured as the midpoint of egg laying plus larval development time. For both treatments, 10 replicates were established with 10 embryos per vial of the experimental diet. Pupae were individually time stamped on the side of the vial every 6 hours during daylight hours.

### Adult weight

Generation 6 adults were collected from development assays, initially weighed for wet weight, then incubated at 55 degrees for 72 hours and weighed again for dry weight.

### RNAseq

Generation 6 embryos were snap frozen in liquid nitrogen. RNA was isolated from ∼20 embryos per cage using the PicoPure RNA isolation kit (Arcturus). DNAse treatment was performed using the RNase-free DNase set (Qiagen). RNAseq was performed at the Ramaciotti centre for genomics (UNSW). RNA was extracted from five biological replicates of each of the 222 and ii2 diet groups (10 samples in total) using the TruSeq Stranded mRNA-seq kit (Illumina, DA, USA). The samples were sequenced in one flowcell on the NextSeq 500 producing paired end reads of 75bp.

## RNA-seq analysis

Reads from the 10 samples were mapped to the NCBI D. melanogaster genome GCF_000001215.4_Release_6_plus_ISO1_MT using the Subread package (v 1.6.3) (Liao et al., 2013). Mapped reads were assigned to features using the featureCounts function of the Subread package, with an average 33.5 M reads assigned per sample. Lowly expressed genes (cpm < 0.5 in at least 4 samples) were filtered out. Differential expression analysis was performed using the Bioconductor packages edgeR (Robinson et al., 2010) and limma (voom) (Ritchie et al., 2015). A DGEList object was created using edgeR and the data was normalised using the TMM method. This DGEList object was used as input in the voom analysis in which the count data was log transformed prior to applying the lmFit and eBayes functions to test for differential expression. Differential expression analysis with edgeR and limma (voom) produced 1550 and 1403 differentially expressed genes (adjusted p.value < 0.05). As 98% of the voom DEGs were also detected by edgeR we used this robust set of genes for downstream analysis, Functional analysis was performed using the kegga and goana funtions of limma, the clustering functions within the STRINGdb package (Franceschini et al., 2012) and the enricher function in the clusterProfiler package (Yu et al., 2012).

### Quantitative PCR

Expression of genes of interest was performed using RT-qPCR. RNA was isolated using Trizol reagent, and cDNA was prepared using Superscript II RTase. Expression of genes of interest were calculated as relative to the average expression of housekeeping genes. Primer sequences for housekeeping genes were: *Act88F* forward 5’-TCGATCATGAAGTGCGACGT-3’, reverse 3’-ACCGATCCAGACGGAGTACT-5’ and *rp49* forward 5’-AGCATACAGGCCCAAGATCG-3’, reverse 3’-TGTTGTCGATACCCTTGGGC-5’.

Primers for the MAPK signalling pathway were: *hkb* forward 5’-TGGCCTTCTCCAACAACG-3’, reverse 3’-CATGTGTTCACGTCGCACTT-5’; *phyl* forward 5’-CGTTCAGCTAATCCAGGCGAA-3’, reverse 3’-GCCTCATTGCTGTTGACCG-5’; *Egfr* forward 5’-CTTAGATAGGAGCCCGTGCG-3’, reverse 3’-CACAAAGATGCAGGGGGACT-5’ and *dsor1* forward 5’-TCAACTGATCGACTCGATGG-3’.

Primer sequences for the ribosome biogenesis pathway were: *nop5* forward 5’-GCCAAGCCGCTGAAAAAGAC-3’, reverse 3’-CTTGATGGCTGTGCCCAGTT-5’; *fib* forward 5’-CTCCGTTGAGACCAATGGC-3’, reverse 3’-ACATGCGAGACTGTCGTTCC-5’; *NHP2* forward 5’-AGTGCTACAAACTGGTGAAGAAG-3’, reverse 3’-GTCGCCAGCAAAGATGCAAA-5’ and *CG3527* forward 5’-AAGGCGATCAATAGGAAGCGT-3’, reverse 3’-CAGCAGCTCGAAAGTGTTGTG-5’.

Primer sequences for the Wnt signalling pathway were: *wntD* forward 5’-TTTGCCATCACATTCTTCATGGG-3’, reverse 3’-GGGTGTACTGGTAGTAGCTCA-5’ and *dnt* forward 5’-ATTGCCACAAGGAACTGCGTTAT-3’, reverse 3’-CCCCAGGCAGTTGTAGTC-5’.

Primers for the Notch signalling pathway were: *DI* forward 5’-TGGAAAGTATTCTCGTCGAAAGC-3’, reverse 3’-TTGTACGGAATGTAGGAGGCG-5’ and *Hey* forward 5’-AGCTGCCAACTGATGAGCC-3’, reverse 3’-GCGAGTCGAGAGTTTTAGACTG-5’.

Primers for the TGF-β signalling pathway were: *Rbf2* forward 5’-GAGACTTGTGAAGTGGAGGGA-3’, reverse 3’-CAAGCGATGGTAGGTGGACAG-5’ and *gbb* forward 5’-GAGTGGCTGGTCAAGTCGAA-3’, reverse 3’-GAAGCCGATCATGAAGGGCT-5’.

Primers for the hedgehog signalling pathway were: *ihog* forward 5’-CGAAGCAATTCCGGTTTTGGA-3’, reverse 3’-TAAGGTACTTGAACCCCGCTC-5’ and *en* forward 5’-GCCAAAGGACAAGACCAACGAC-3’, reverse 3’-CGGTCAGATAGCGATTCT-5’.

Primers for ribosome copy number were *tRNAK-CTT* forward 5’-CTAGCTCAGTCGGTAGAGCATGA-3’, reverse 3’-CCAACGTGGGGCTCGAAC-5’ and *18srDNA* forward 5’-AGCCTGAGAAACGGCTACCA-3’, reverse 3’-AGCTGGGAGTGGGTAATTTAC.

### Expression at different ages and tissues

Flies were assessed for expression of MAPK and Ribosome biogenesis pathway genes using RT-qPCR at 4 life stages: egg, and in whole body and isolated fat bodies from 2^nd^ instar larvae, 5 d adults and 15 d adults.

### Statistics

All data were analysed for normality by Shapiro-Wilks W test to remove outliers. Outliers were identified as values greater than 1.5 times the interquartile range. ANOVA analyses were conducted for the development assays with the main effect of ancestral diet, while expression at different ages and tissues were analysed with main effects of age, ancestral diet, and their interaction. For all other biochemical assays, tow-tailed Student’s t-tests were conducted.

## Results

### Development time

The 222 flies developed significantly faster than all other treatments, with ii2 and 442 having the same rate of development, and 662 having the slowest (Fig. 2A, B). Ancestral diet significantly affected time to pupation (F_3, 365_ = 105.18, p < 0.0001) and 3 d eclosion (F_3, 36_ = 225.69, p < 0.0001). The effects on developmental timing were also seen when a single previous generation was exposed to a particular diet (Fig. S1). This indicated that the ancestral diet effect can be induced by a single generation, rather than being accumulative or exacerbated by an increased number of exposures. Adult dry-weight was measured to determine if the faster development of the 222 flies was perhaps a trade-off for lower body size, which is correlated with adult fecundity (Yadav and Sharma, 2014). No difference in body weight was observed (t_18_ = 0.17, p = 0.87, data not shown).

**Figure 2.**
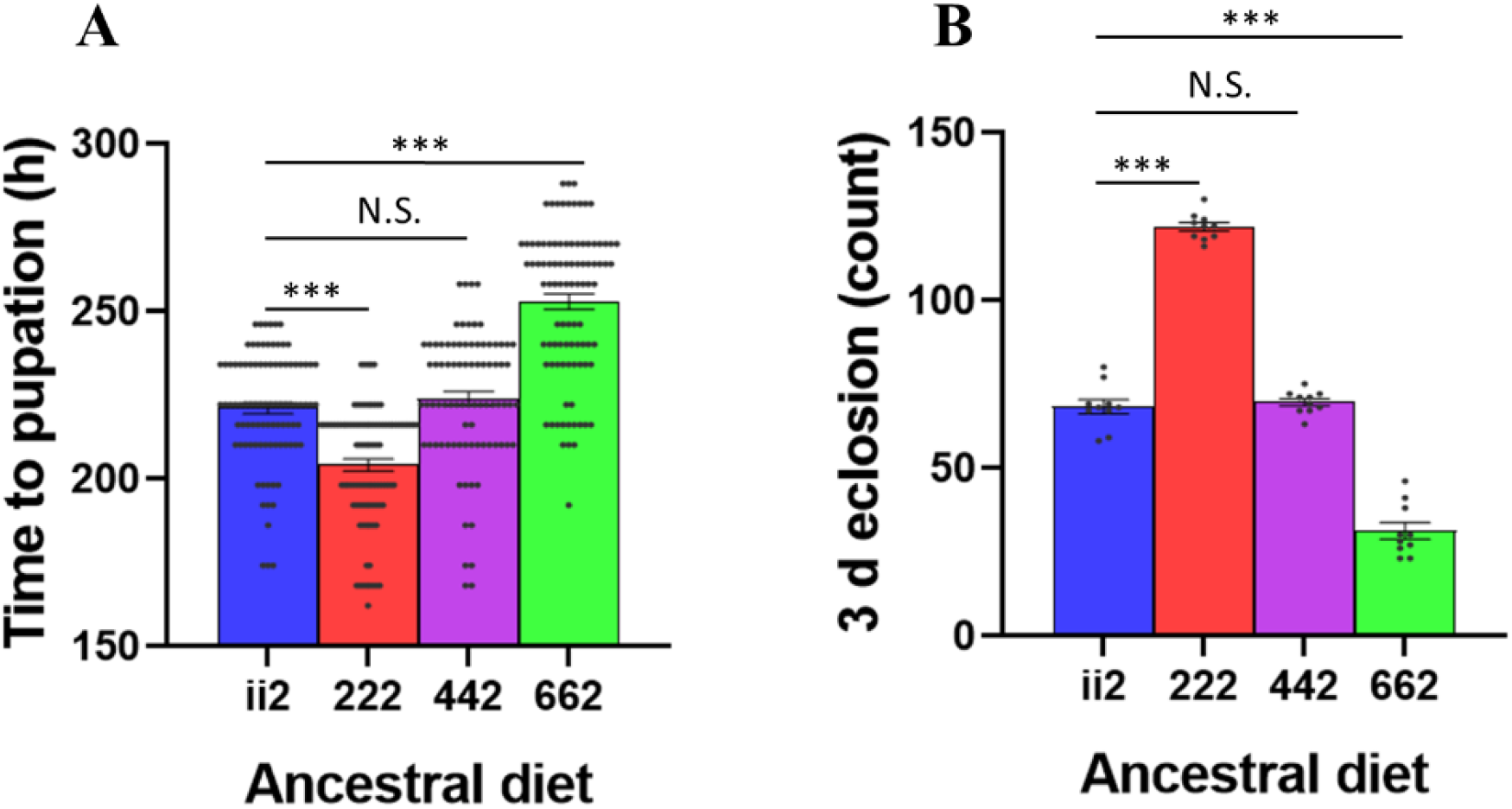
Development assays: Time to pupation (A) and 3 d eclosion (B) of the 222 flies was faster than that of the ii2 and 442 flies, while the 662 flies were the slowest to develop. Bars indicate mean values ± SEM. N.S. indicates not significant, *** indicates p < 0.001.

### Transcriptome analysis of embryos

The ancestral diet-induced differences in developmental timing were evident in the period between embryonic and pupal stages. Therefore, we performed RNA-seq on ii2 and 222 embryos to identify formative transcriptional signatures that could drive the differences in growth timing. A multidimensional scaling plot demonstrated that the 5 replicates of the 222 flies separated from the 5 replicates of the ii2 flies along the main principal component consistent with a different transcriptional signature in these conditions (Fig 3A). Differential expression analysis with limma (voom) produced 1403 differentially expressed genes (DEGs) (adjusted p.value < 0.05). 851 of the voom DEGs were upregulated and 552 were downregulated in the 222 samples compared to ii2 as illustrated by the volcano plot (Fig 3B). Pathway and network analysis of the DEGs revealed that growth-associated pathways such as ribosome biogenesis, Wnt signaling and MAPK signaling were higher in 222 compared to ii2 (Fig. 3C-E).

**Figure 3.**
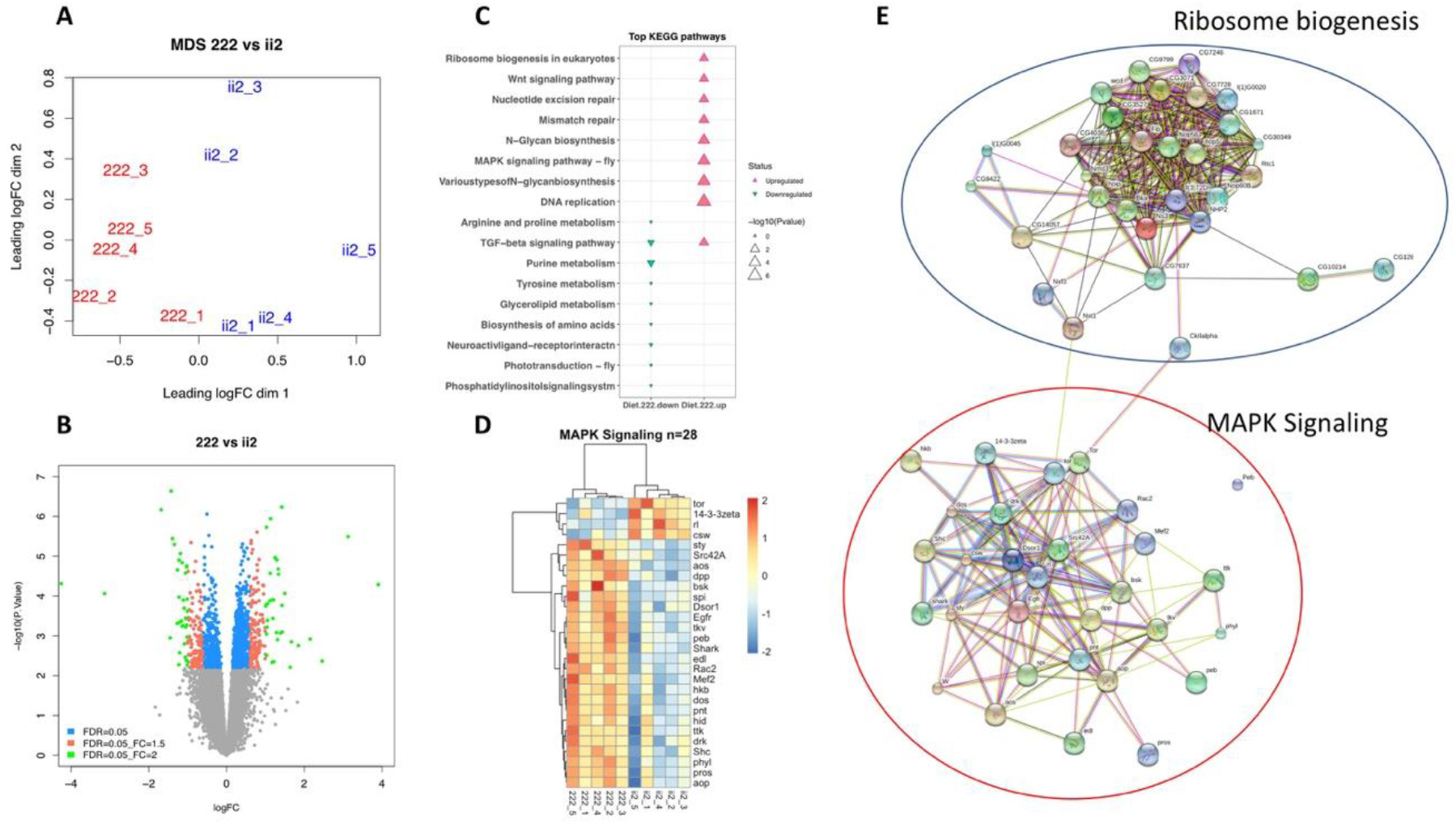
RNAseq analysis: (A) Multi dimensional scaling plot showing separation between samples (222 = 3 generations fed on 1:2 Protein: Carbohydrate diet, ii2= change in ancestral diet). (B) Volcano plot showing gene expression changes associated with the 222 diet. (C) Top KEGG pathways as found using kegga (limma). (D) heatmap of DEGs involved in MAPK Signaling pathway. E. Differentially expressed genes between the 222 diet and the ii2 diet involved in the Ribosome and MAPK signaling pathways with a FDR < 0.1.

### Pathway validation

Expression of the top two genes from each of the identified development pathways was measured. MAPK and Ribosome Biogenesis were identified as the top two pathways, while Hedgehog did not pass validation (Fig. 4).

**Figure 4.**
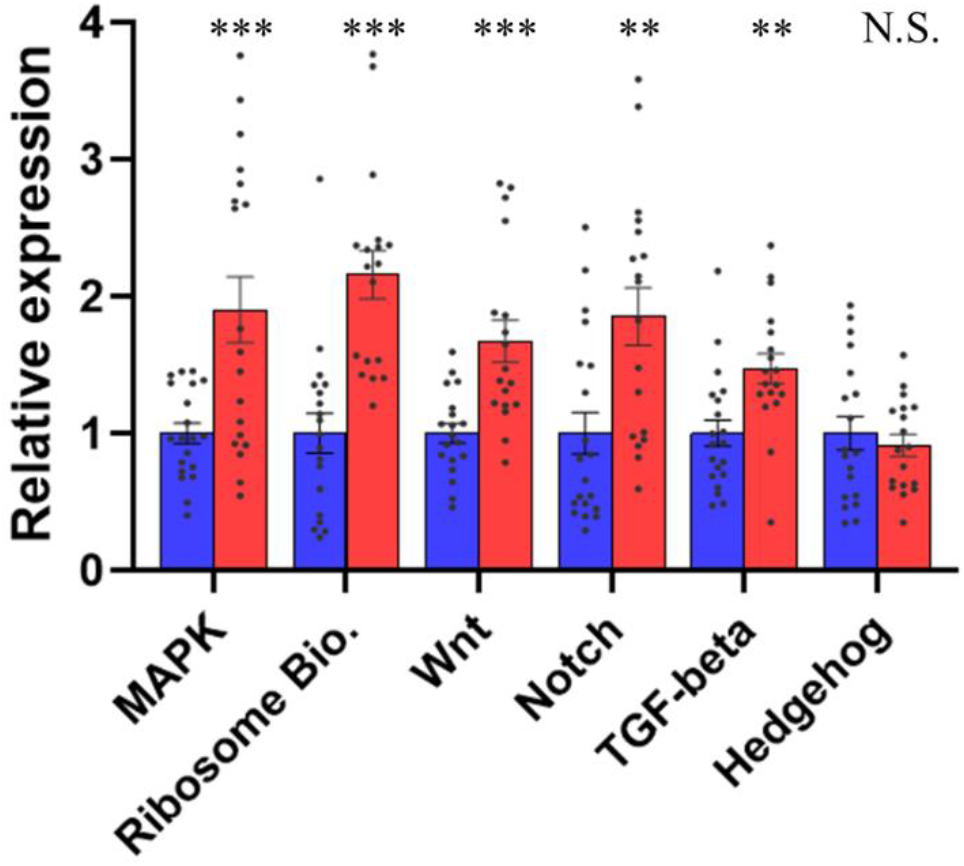
Pathway validation: MAPK, Ribosome Biogenesis, Wnt, Notch and TGF-beta passed validation (t_37_ = 3.67, p <0.001; t_35_ = 5.54, p < 0.001; t_35_ = 3.82, p < 0.001; t_37_ = 3.06, p = 0.004; t_36_ = 3.22, p = 0.003, respectively), while hedgehog did not (t_35_ = 0.32, p = 0.75). MAPK genes *hkb* and *phyl* (t_18_ = 2.38, p = 0.03; t_17_ = 2.69, t = 0.02, respectively), Ribosome Biogenesis genes *nop5* and *Fib* (t_17_ = 7.21, p < 0.001; t_16_ = 2.11, p = 0.05, respectively), Wnt genes *WntD* and *dnt* (t_16_ = 3.20, p = 0.006; t_17_ = 2.71, p = 0.01, respectively), Notch genes *DI* and *Hey* (t_17_ = 1.45, p = 0.17; t_17_ = 3.37, p = 0.004, respectively), TGF-beta genes *Rbf2* and *gbb* (t_17_ = 2.55, p = 0.02; t_17_ = 1.96, p = 0.07, respectively), and Hedgehog genes *ihog* and *en* (t_16_ = 2.04, p = 0.06; t_17_ = 2.65, p = 0.02, respectively). Bars indicate pooled relative expression ± SEM. Blue bars indicate ii2 diet, red bars indicate 222 diet. ** indicates p < 0.01, *** indicates p < 0.001.

Two further genes were identified from each pathway for analysis of the top two pathways. For MAPK signalling genes (Fig. S2A), expression was higher in 222 for *hkb, phyl, Egfr*, and *Dsor1* (t_17_ = 7.81, p < 0.001; t_15_ = 3.09, p = 0.008; t_14_ = 2.47, p = 0.03; t_16_ = 2.44, p = 0.03, respectively). For Ribosome biogenesis pathway genes (Fig. S2B), expression was higher in 222 for *nop5, Fib, NHP2*, and *CG3527* (t_18_ = 2.38, p = 0.03; t_17_ = 2.69, p = 0.02; t_15_ = 2.23, p = 0.04; t_15_ = 2.27, p = 0.04, respectively).

### Expression at different ages and tissues

To determine whether this was just an early developmental effect, expression of the four identified genes for MAPK signalling and Ribosome biogenesis were assayed from RNA extracted from eggs, as well as from the whole body of larvae, 5 d adults and 15 d adults (Fig. 5) and from fat bodies of larvae, 5 d adults and 15 d adults (Fig. S3).

**Figure 5.**
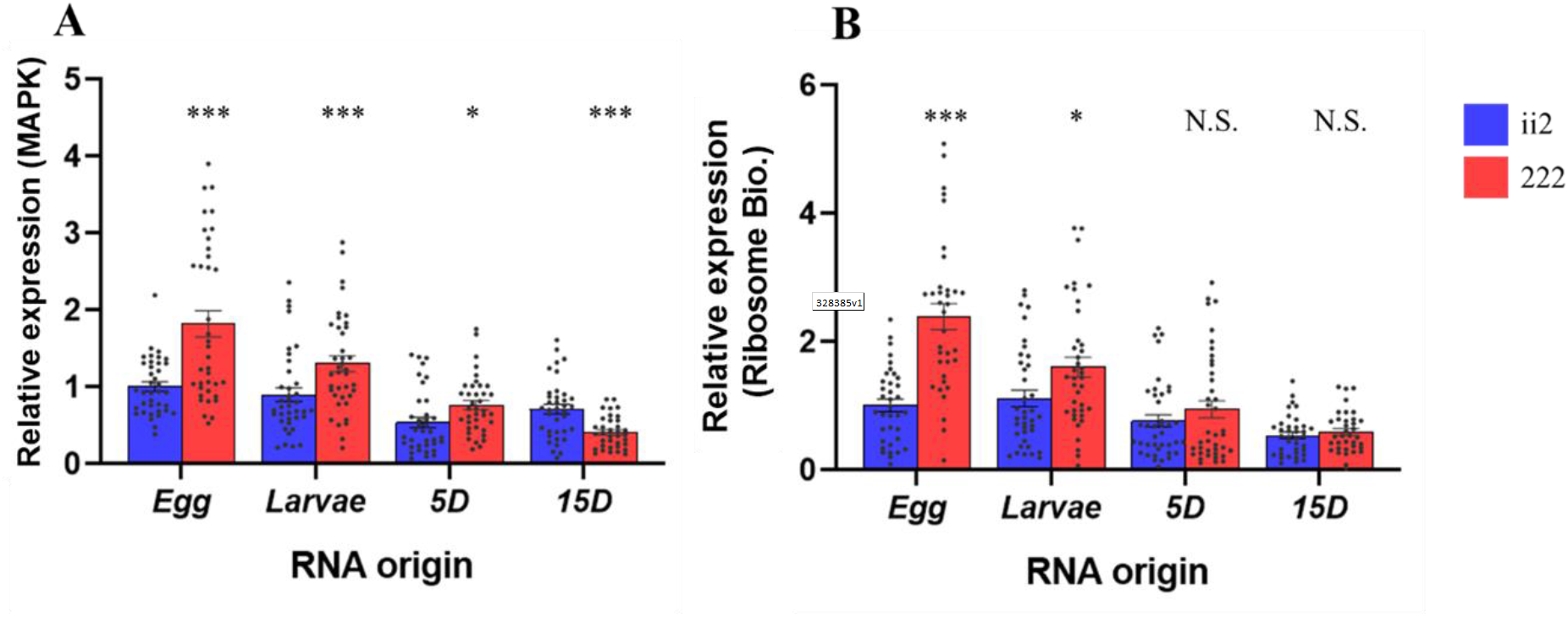
Expression at different life history stages: Expression of MAPK genes (A) and of Ribosome genes (B) decreased with age. Expression of MAPK was different between ii2 and 222 in eggs and larvae, while ribosome gene expression was only different in eggs. Bars indicate relative expression ± SEM. Bars not connected by the same letter differ significantly according to Tukey’s HSD test.

For MAPK genes ANOVA identified a significant effect of Age, Diet, and the interaction of Age with Diet (F_3, 292_ = 39.74, p < 0.001; F_1, 292_ = 20.40, p < 0.001; F_3, 292_ = 13.25, p < 0.001, respectively), with expression decreasing with age in both whole body and fat bodies (Fig. 5A, Fig. S5A).

For Ribosome biogenesis genes, ANOVA identified a significant effect of age, diet and age by diet interaction (F_3, 292_ = 32.74, p < 0.001; F_1, 292_ = 36.43, p < 0.001; F_3, 292_ = 11.40, p < 0.001, respectively) with expression decreasing with age in both whole body and fat bodies (Fig. 5B, Fig. S5B).

### Phenocopy of ancestral-diet induced effects on developmental timing with pharmacological inhibition of MAPK and ribosome biogenesis pathways

Chemical inhibition of developmental pathways can lead to morphological differences resulting in a copy of a phenotype (phenocopy) that is distinct to their normal phenotype (Aw et al., 2018). We partially inhibited the MAPK and ribosome biogenesis pathways of the 222 flies to attempt to phenocopy the slower ii2 rate of development (Fig. 6).

**Figure 6.**
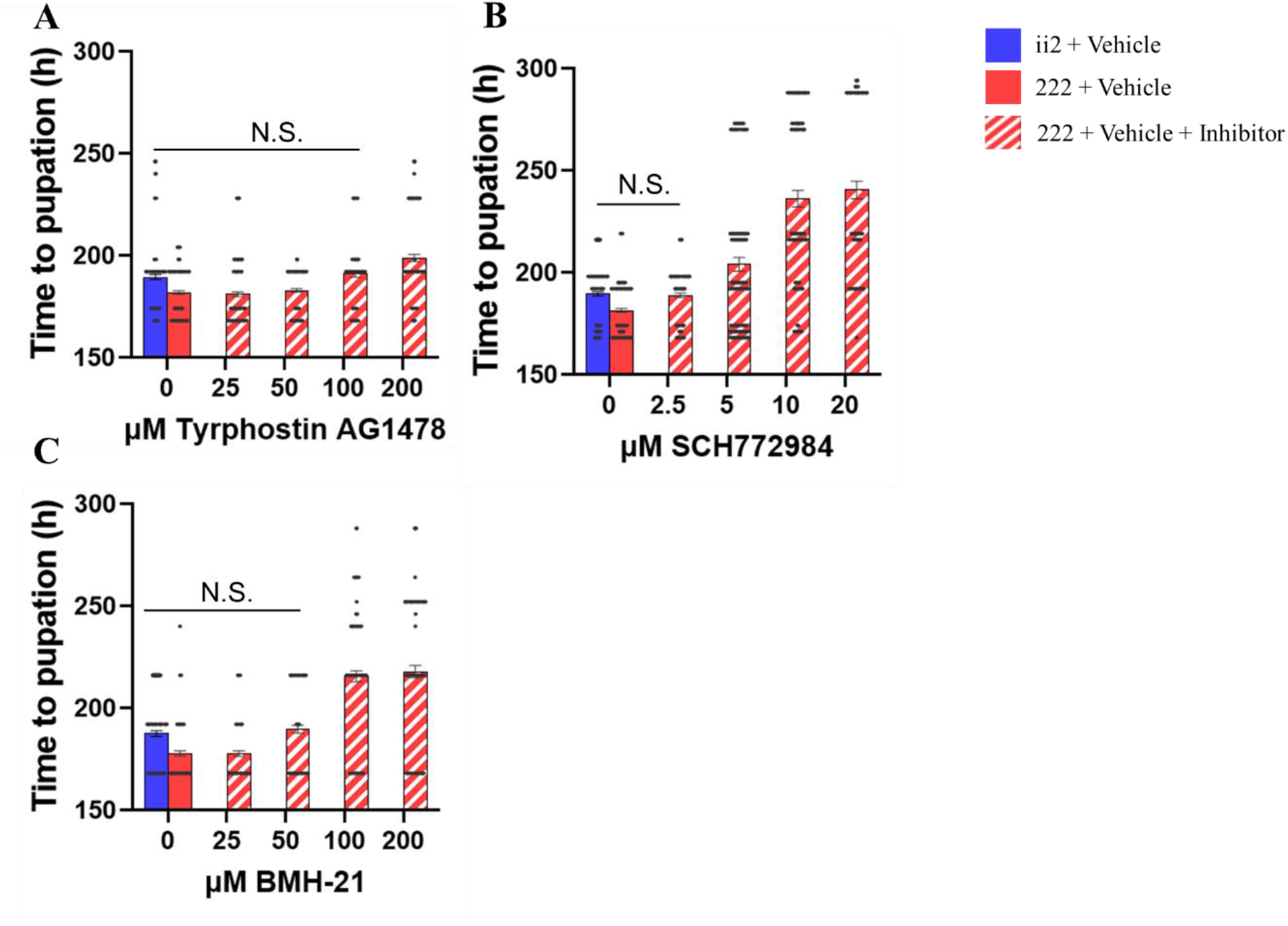
Pathway inhibitors: Chemical inhibition of the MAPK and Ribosome Biogenesis pathways in 222 flies resulted in phenocopies of the ii2 phenotype. Phenocopies of ii2 were observed when fed (A) 100 μM Tyrphostin AG1478, an EGFR inhibitor, (B) 2.5 μM of SCH772984, an ERK 1/2 inhibitor, and (C) 50 μM of BMH-21, a RNA polymerase I inhibitor.

The EGFR inhibitor Tyrphostin AG1478 slowed development of 222 at 100 μM concentration to the same rate as ii2, with treatment having a significant effect (F_5, 1001_ = 28.91, p < 0.001) (Fig. 6A). The ERK1/2 inhibitor SCH772984 slowed development of 222 to be the same as ii2 at 2.5 μM concentration with treatment having a significant effect (F_5, 880_ = 128.73, p < 0.001) (Fig. 6B). The RNA polymerase I inhibitor BMH-21 slowed development of 222 flies to be the same as ii2 at 50 μM concentration, with treatment having a significant effect (F_5, 874_ = 77.20, p < 0.001) (Fig. 6C).

### Gene expression analysis of pharmacological inhibition of pathways

Partially blocking MAPK at EGFR with 100 μM Tyrphostin AG1478 in 222 flies showed reduced expression of genes in the MAPK pathway (t_17_ = 2.80, p = 0.01; t_18_ = 2.82, p = 0.01; t_18_ = 2.24, p = 0.04; t_18_ = 3.32, p = 0.004 for *hkb, phyl, Egfr*, and *Dsor1*, respectively, Fig. 7A), and Ribosome biogenesis pathways (t_18_ = 3.28, p = 0.005; t_18_ = 5.41, p < 0.001; t_18_= 3.51, p = 0.003; t_18_ = 3.29, 0 = 0.004 for *nop5, Fib, NHP2*, and *CG3527*, respectively; Fig. 7B).

**Figure 7.**
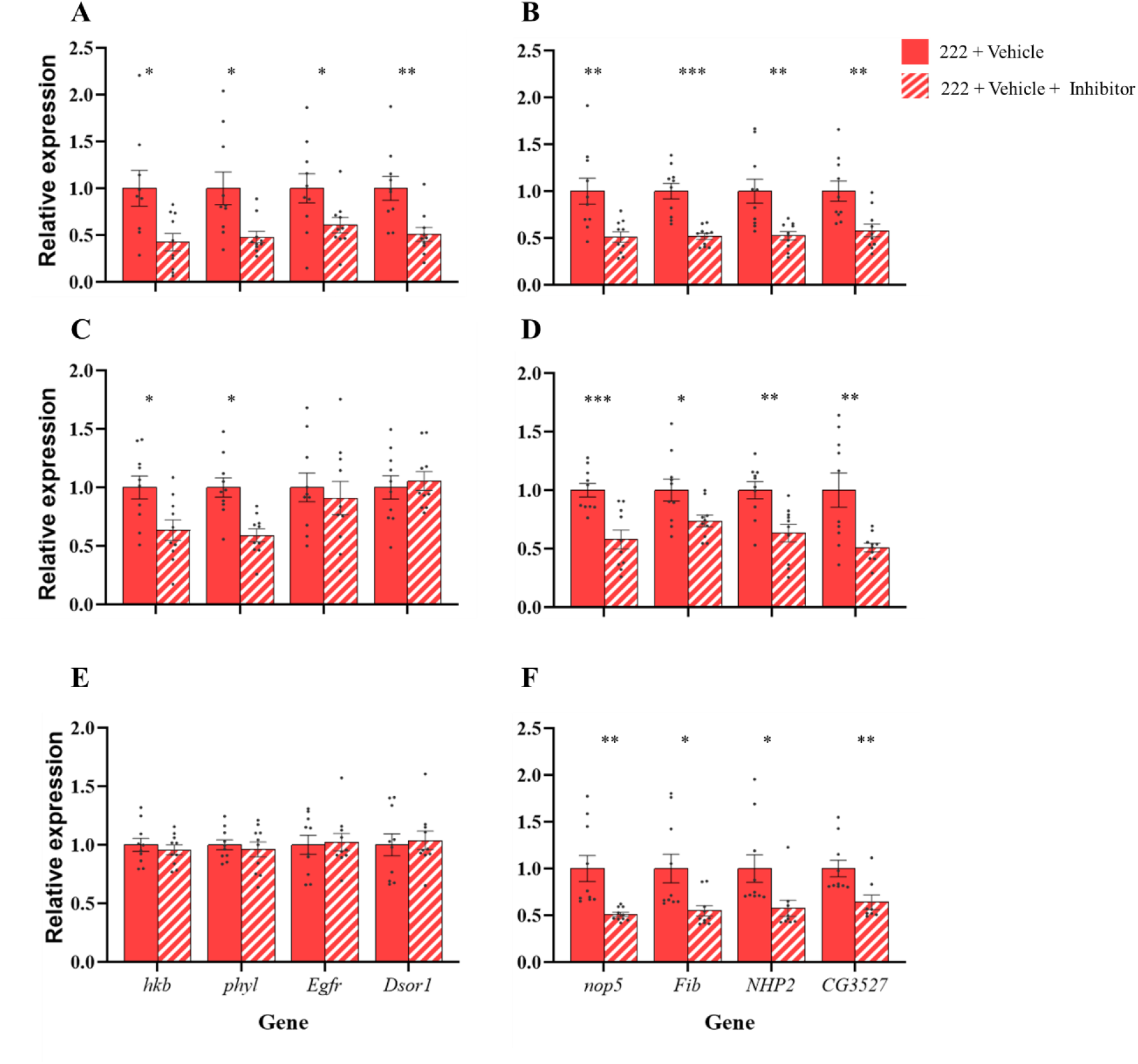
Inhibited pathway qPCR: Blocking of MAPK with 100 μM Tyrphostin AG1478 or 2.5 μM of SCH772984, showed a reduction of MAPK gene expression at the point of inhibition and downstream (A, C) but also led to a reduction in expression of ribosome biogenesis pathway genes (B, D). Partially inhibiting ribosome biogenesis with 50 μM of BMH-21 showed no change in MAPK expression (E) but did show a reduction in ribosome biogenesis pathway genes (F). Bars indicate relative expression ± SEM. * indicates p < 0.05, ** indicates p < 0.01, *** indicates p < 0.001.

Partial blocking of MAPK at ERK1/2 with 2.5 μM SCH772984in 222 flies showed reduced expression of MAPK genes downstream of the inhibitor, with no difference in expression upstream (t_18_ = 2.77, p = 0.01, t_18_ = 4.14, p = 0 < 0.001, t_18_ = 0.48, p = 0.63, t_18_ = 0.43, p = 0.67 for *hkb, phyl, Egfr*, and *Dsor1*, respectively, Fig. 7C). Ribosome biogenesis pathways were also down-regulated (t_18_ = 4.28, p < 0.001, t_18_ = 2.47, p = 0.02, t_18_ = 3.51, p = 0.003, t_18_ = 3.27, p = 0.004 for *nop5, Fib, NHP2*, and *CG3527*, respectively; Fig. 7D).

However, partial inhibition of ribosome biogenesis with 50 μM of BMH-21 showed no significant difference in MAPK pathway genes (t_18_ = 0.61, p = 0.55; t_18_ = 0.51, p = 0.61; t_18_ = 0.21, p = 0.84; t_18_ = 0.31, p = 0.76 for *hkb, phyl, Egfr*, and *Dsor1*, respectively, Fig. 7E), but did show reduced expression of ribosome biogenesis pathway genes (t_18_ = 3.48, p = 0.003; t_18_ = 2.78, p = 0.01; t_17_ = 2.43, p = 0.03; t_16_ = 2.96, p = 0.009 for *nop5, Fib, NHP2*, and *CG3527*, respectively; Fig. 7F).

## Discussion

Here we have shown in a new *Drosophila* model that the MAPK pathway is a mediator of grand-parental dietary P:C ratio effects on prepupal developmental timing. The overall effect of a higher ancestral P:C ratio (e.g. P:C 1:2) is faster development through the embryo to pupal stages than a lower ratio (e.g. P:C 1:6). This result was highly reproducible, seen in over eight independent experiments, and induced either when the grand-parental, or the grand-parental plus great grand-parental generation had the diets.

The study design did not replicate our previous investigations of multiple generational dietary exposure (Aw et al., 2018). The dietary formulations, and use of the Alstonville strain were the same, but there were key differences such as the number of generations that were exposed to a particular diet, and the number of fly strains used. It was essential to change the study design to remove the influence of parental diet on offspring development which have been well described (Emborski and Mikheyev, 2019; Guida et al., 2019; Matzkin et al., 2013; Strilbytska et al., 2020; Valtonen et al., 2012; Vijendravarma et al., 2010; Zajitschek et al., 2017).

In comparison with our previous work, the close similarity of the effects of a two-generation and one-generation dietary exposure (Suppl Figure) suggests that the new model is not reproducing the accumulative transgenerational effects (Aw et al., 2018). This may be due to the accumulative effect occurring after a larger number of generations (it was previously seen after 4 generations on a diet), or it could be linked to inter-strain competition as it was previously seen in flies kept in cages containing 2-4 different *Drosophila* strains.

Diet is a strong determinant of developmental timing in animals (Danielsen et al., 2013; Nagarajan and Grewal, 2014). In *Drosophila* the effects of varying carbohydrate, protein and lipid on the timing of the various developmental stages have been examined, and the importance of the insulin/TOR pathway for regulation has been highlighted (Danielsen et al., 2013). In this pathway circulating levels of amino acids, carbohydrates and *Drosophila* insulin-like peptides are detected by cell surface receptors which activate intracellular networks of signalling molecules including PI3K/AKT, FOXO and TOR which in turn regulate processes required for growth such as transcription, protein synthesis and autophagy (Danielsen et al., 2013). However, it is important to note that in our study the developmental timing changes in offspring were due to their ancestors’, not their own diet, thus the flies on the ii2 or 222 diets will have qualitatively equivalent circulating levels of amino acids and carbohydrates. Indeed, the RNA-seq analysis did not indicate that this key growth regulatory pathway is responsible for the observed ancestral diet effects.

In invertebrates, levels of dietary protein in particular have been found to contribute to growth, fecundity and ageing (Auge et al., 2017; Bruce et al., 2013; Lee et al., 2008a). Faster developmental timing is observed when a high P:C ratio is consumed by larvae (Aw et al., 2018), by only the parental generation (Matzkin et al., 2013) and now in this study, only in the grand-parental generation. A recent study of the effects of an ancestral ‘rich’ relatively high protein vs ‘poor’ lower protein diets also observed a faster developmental timing in grand-offspring of the ‘rich’ diet flies (Deas et al., 2019). Interestingly, they observed a greater influence of grand-parental diet than parental diet on a variety of offspring phenotypes, including larval to pupal developmental timing. However, the study did not investigate the underlying growth regulatory pathways that are affected by the different generational dietary exposures diet.

Progression to the next larval stage in Drosophila is triggered by the larva attaining a critical body weight, which leads to activation of a MAPK signalling cascade which upregulates the expression of ecdysone biosynthetic genes (Rewitz et al., 2009; Tennessen and Thummel, 2011). Our examination of embryo transcriptome suggests that is possible that alterations to this pathway may explain the observed differences in developmental timing due to ancestral diet. However, in addition to ecdysone biosynthetic genes, the downstream targets of altered MAPK signalling include those responsible for ribosomal RNA biogenesis which is a crucial for generation of proteins and thus cellular growth (Sriskanthadevan-Pirahas et al., 2018). We were able to phenocopy the ancestral diet-induced effects on developmental timing through pharmacological inhibition of both MAPK proteins ERK1/2 and EGFR as well as ribosomal biogenesis. These experiments also confirmed the hierarchy of the 2 processes as inhibition of MAPK proteins affected rRNA transcript levels but not *vice versa*.

Another MAPK regulated process that could explains the changes in developmental timing is appetite and food intake. We did not observe a difference in the adult offspring dry body weight which suggests that that there was no trade-off between growth timing and eventual body size. This also indicates that larval size was not different either and thus the faster developing flies were able to achieve the equivalent critical body weight at each larval stage. To achieve this the larvae with ancestral P:C 1:2 exposure would have to eat faster than those with P:C 1:6 diet ancestors. This may be facilitated through programming of olfactory systems as a gene with one of the largest expression fold increases in 222 compared to ii2 was *Gustatory receptor 64f (Gr64f)*. This gene is expressed in gustatory receptor neurons and detects sugars suggesting that these flies have an increased ability to detect food cues (Branch and Shen, 2017; Jiao et al., 2008). Furthermore, in *Drosophila* and mouse, MAPK signalling has been shown to be an important part of appetite regulation (Lee et al., 2008b; Rodrigues et al., 2017). Therefore, our study may provide an example of ancestral P:C ratio influencing offspring and grand-offspring food intake. Intriguingly, a recent mouse study also showed that a MAPK pathway is altered in the brains of offspring from obese mothers, which the authors proposed to contribute to altered offspring appetite regulation and subsequent risk of obesity (Bae-Gartz et al., 2019). In our study differences in the MAPK genes which had altered transcript levels at the embryo stage were also seen at different life stages and even adult tissues which suggests that programming of MAPK-regulated processes could be pervasive and include those involved in appetite regulation.

Finally, while providing new insight into how transgenerational effects are mediated in the post-fertilisation period, our study does not provide evidence on what the molecular change is in the gametes that drives the subsequent developmental changes. A previous transgenerational study in *Drosophila* found rDNA copy number reduction in offspring of flies that consumed a high protein diet (Aldrich and Maggert, 2015). We found no evidence for this genetic change in the offspring in our study which points to an epigenetic source of the growth phenotype. The prominence of the MAPK pathway in the network analysis of embryonic transcriptomes in our study make it tempting to predict that the transgenerational phenotypes are caused by inheritance of histone modifications or small non-coding RNA that modulates this pathway.

## Acknowledgements

This study was funded by an Australian Research Centre Discovery Project Grant (DP190102555).

The RNA-seq bioinformatics analysis was performed by SC in the Systems Biology Initiative UNSW headed by Professor Marc Wilkins who is thanked for his assistance. SC’s work within the SBI was funded by the Australian Federal Government CRIS scheme and a UNSW RIS grant.

## Supplementary figures

**Supplemntary Figure 1.**
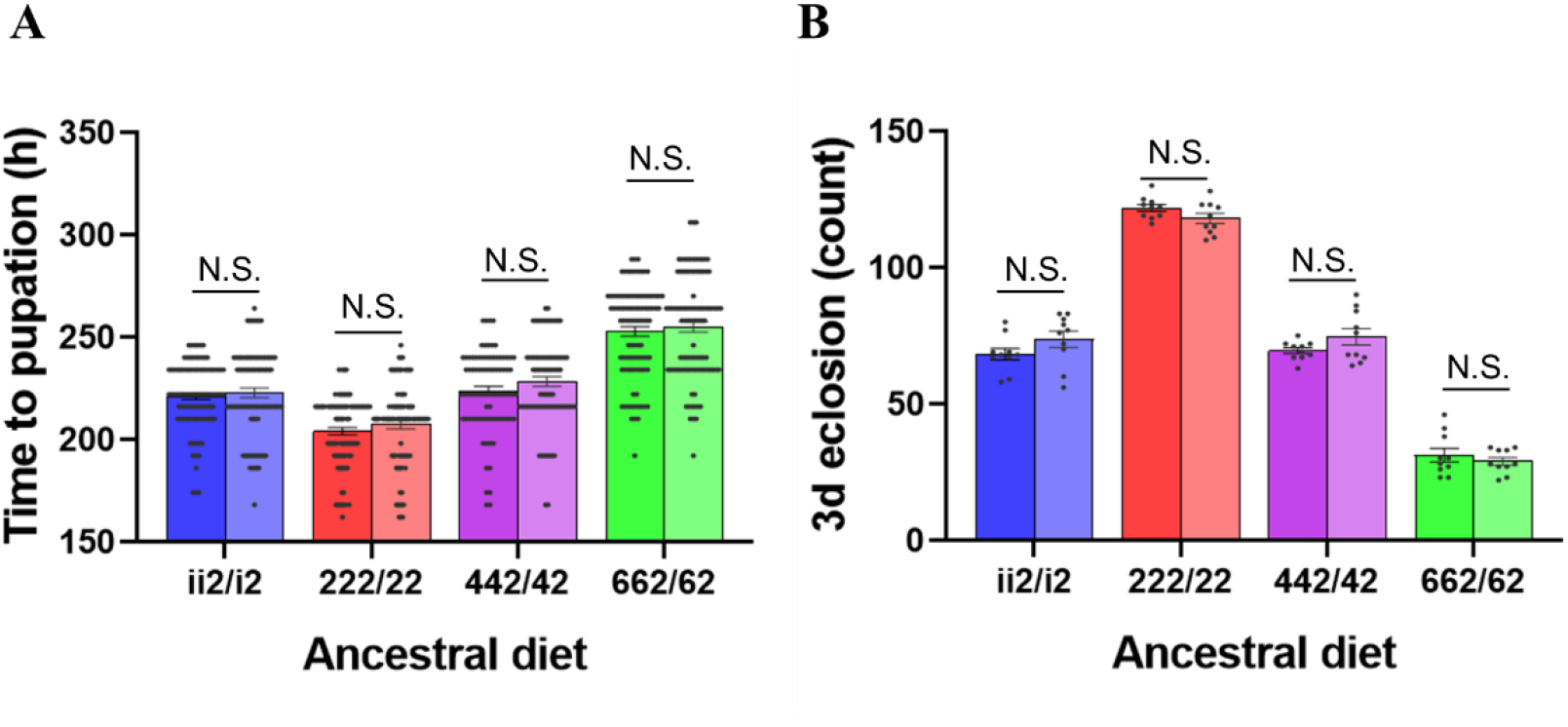
Development assays one or two generations: providing a different ancestral diet for one or two generations did not signifcantly effect rate of development. Time to pupation showed a significat effect of ancestral diet, but no effect of single/double generation of the ancestral diet or the interaction (F_3, 724_ = 179.09, p < 0.001; F_1, 724_ = 3.68, p = 0.06; F_3, 724_ = 0.16, p = 0.92, respectively). 3d eclosion was also unaffected by providing a different ancestral diet for one or two generations. 3d eclosion showed asigificant effect of diet, but no effect of single/double generation of ancestral diet or the interaction (F_3, 72_ = 584.47, p < 0.001; F_1, 72_ = 0.52, p = 0.47; F_3, 72_ = 2.60, p = 0.06, respectively). Bars indicate mean values ± SEM.

**Supplementary figure 2.**
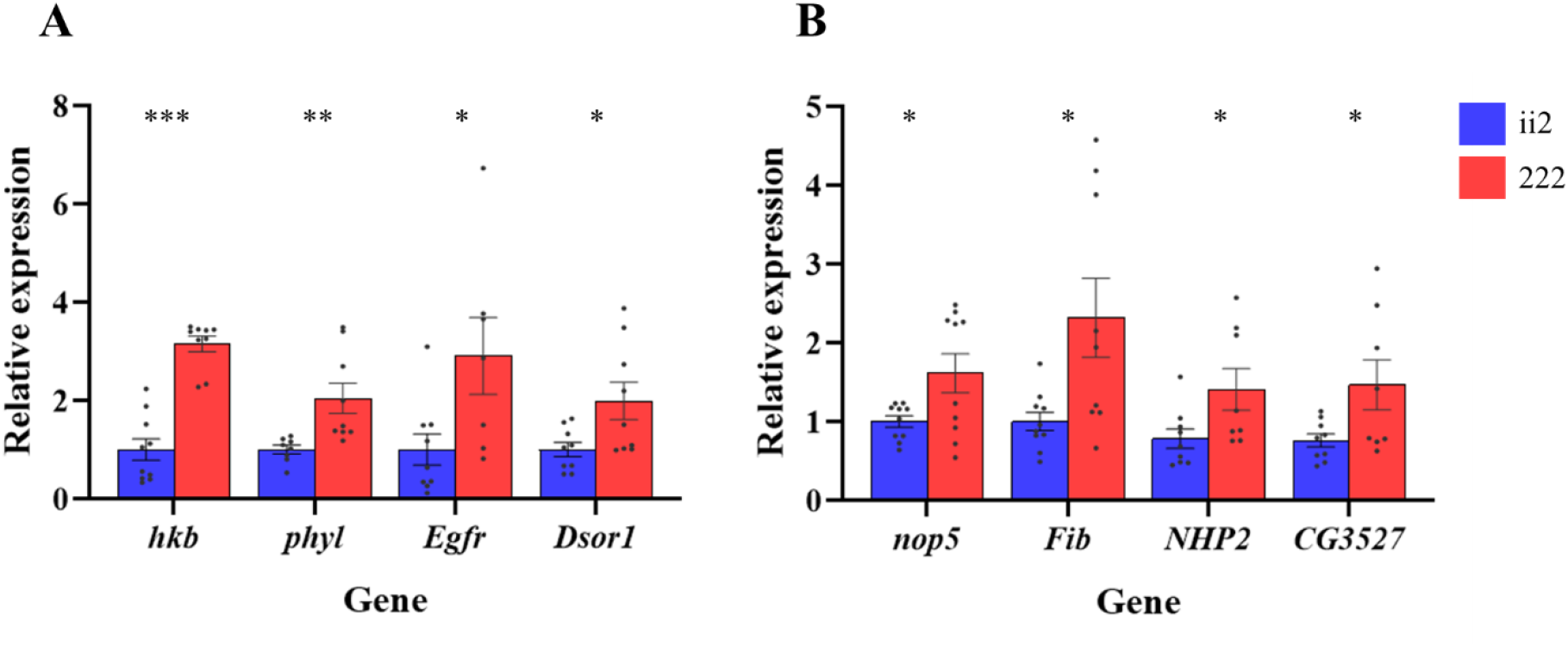
Top pathway validation: Expression of MAPK signalling pathway genes (A) and Ribosome biogenesis genes (B) was higher for the 222 flies. Bars indicate relative expression ± SEM. * indicates p < 0.05, ** indicates p < 0.01, *** indicates p < 0.001.

**Supplementary Figure 3.**
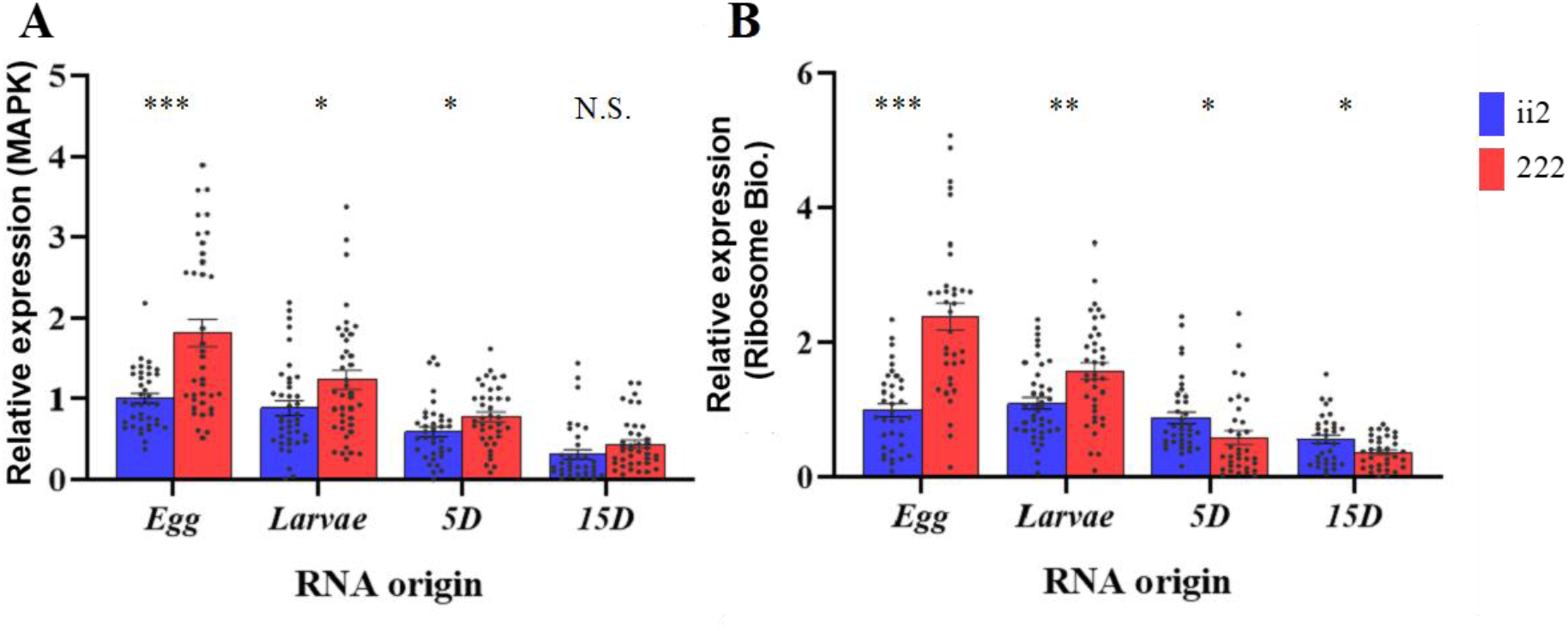
Expression at different ages (Fat body): The ii2 treatment had significantly higher expression for the MAPK pathway (A) as eggs (t_73_ = 4.58, p < 0.001) as well as from larval fat bodies (t_78_ = 2.37, p = 0.02), 5 day old adult (5D) fat bodies (t_72_ = 2.16, p = 0.03;), but not 15 day old adult (15D) fat bodies (t_72_ = 1.66, p = 0.10). For the Ribosome biogenesis pathway (B) ii2 eggs had significantly higher expression (t_69_ = 6.24, p < 0.001), as did larval fat bodies (t_76_ = 3.11, p = 0.003), 5 day old adult fat bodies (t_69_ = 2.19, p = 0.03). In 15D adult fat bodies the 222 treatment had significantly higher expression (t _66_ = 2.65, p = 0.01).

## Notes

### Competing Interest Statement

The authors have declared no competing interest.

